# Towards an emotional stress test: a reliable, non-subjective cognitive measure of anxious responding

**DOI:** 10.1101/062661

**Authors:** Jessica Aylward, Oliver J Robinson

**Affiliations:** UCL Institute of Cognitive Neuroscience

## Abstract

Response to stress is a key factor in mood and anxiety disorder aetiology. Current measures of stress-response are limited because they largely rely on retrospective self-report. Objectively quantifying individual differences in stress response would be a valuable step towards improving our understanding of disorder vulnerability. Our goal is to develop a reliable, objective, within-subject probe of stress response. To this end, we examined stress-potentiated performance on an inhibitory control task from baseline to 2-4 weeks (n=50) and again after 5-9 months (n=22) as well as examining population measures for a larger sample (n=165). Replicating previous findings, threat of shock improved distractor accuracy and slowed target reaction time on this task. Critically, both within-subject self-report measures of stress (ICC=0.74) and stress-potentiated task performance (ICC=0.58) showed clinically useful test-retest reliability. Threat-potentiated task performance may therefore hold promise as a non-subjective measure of individual stress-reactivity.

## Introduction

Mood disorders are common, but there is huge variability in vulnerability (Kendler, Kuhn, & Prescott, 2004). According to the diathesis–stress model (Monroe & Simons, 1991), a disorder is triggered when an underlying vulnerability, coupled with stressful life events reaches a threshold. This suggests that a clear ability to quantify mood and anxiety disorder vulnerability will not emerge without an ability to quantify individual differences in stress response.

Stress induced experimentally using threat of unpredictable shock engages similar circuitry as pathological anxiety (Robinson, Letkiewicz, Overstreet, Ernst, & Grillon, 2011). In this paradigm, unpredictable electric shocks are delivered to the wrist, independent of task performance. The impact of stress is investigated by comparing performance in the same individual when they are at risk and safe from shock. Many domains of cognition are affected (for a review see Robinson, Vytal, Cornwell, & Grillon, 2013) including inhibitory control (Robinson, Vytal, et al., 2013) and attentional bias towards threat (Cornwell et al., 2011).

The Sustained Attention to Response Task (SART), where participants respond to frequent target stimuli whilst withholding a response to infrequent distractor stimuli (under alternating threat and safe conditions), can explore the interaction between stress and inhibitory control. This impact of stress on this task has been confirmed in numerous studies – threat of shock improves accuracy and slows down responses (Aylward, Roiser, & Robinson, 2015; Grillon, Robinson, Mathur, & Ernst, 2016a; Mkrtchian, Roiser, & Robinson, 2015; Robinson, Krimsky, & Grillon, 2013) – but its test-retest reliability is unknown.

In the cardiac stress test, subjecting the heart to stress reveals key diagnostic signatures of heart disease vulnerability that are not evident at rest (Peteiro, 2010). Here we seek to develop an emotional equivalent of the cardiac stress test. However, in order to have clinical value, test performance needs to be reliable and stable across time in the same individuals (Alonso, Geys, Molenberghs, & Vangeneugden, 2002). Many cognitive tasks, nevertheless, show poor reliability (see Table 1).

**Table 1:**
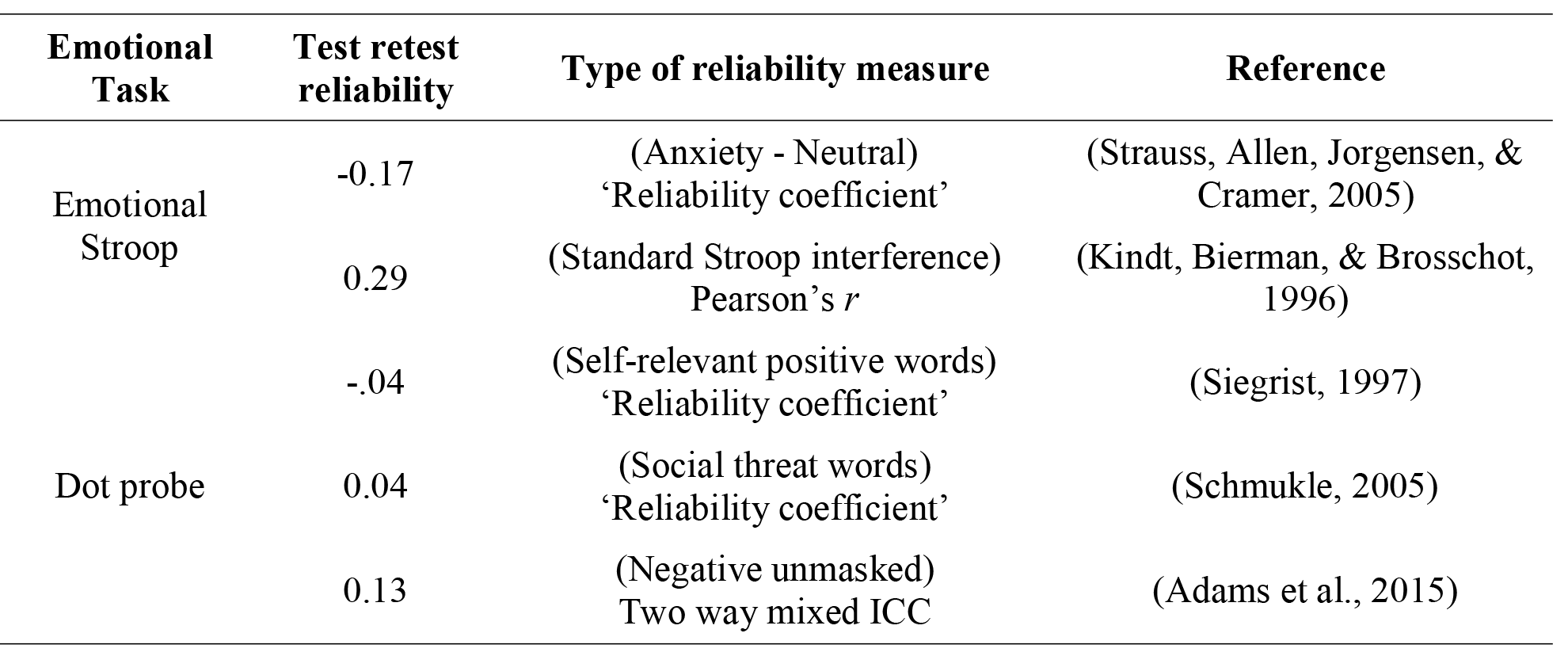
*Commonly used emotional tasks and their test re-test reliabilities. Note that for the reliability coefficients / Pearsons’s *r*: 0.7 is strong, 0.5 is moderate and 0.3 is weak reliability For the ICCs, 0.4-0.75 is ‘fair to good’ reliability and >0.75 is ‘excellent’ reliability*.

Here we therefore probed the test-retest reliability of stress responding on the SART using Intraclass Correlation Coefficients (Fleiss, 1986). We predicted that threat of shock would improve accuracy at withholding a response to distractor stimuli and slow down responding to target stimuli, in line with previous findings. Critically, we predicted that this would be reliable across testing sessions and hence constitute a non-subjective, behavioural measure of individual stress response.

## Method

Fifty healthy participants (25 female, mean age = 26.5, SD = 8.47), completed the SART in two testing sessions, separated by a period of between two and four weeks. Twenty two participants (11 female, mean age = 28.5, SD = 11.00) completed the task for the third time in a follow up session between five and nine months later. A screening procedure prior to participation verified that participants had no history of neurological, psychiatric, or cardiovascular conditions. Exclusion criteria also included alcohol dependence and any recreational drug use in the last 4 weeks.

The methods were identical on each session. Participants provided written informed consent to take part in the study (UCL ethics reference: 1764/001). Prior to participation, subjects were screened to ensure that they had no history of neurological, psychiatric, or cardiovascular conditions.

An *a priori* power analysis was run in G*Power (Faul, Erdfelder, Lang, & Buchner, 2007). The power analysis was based on previous results of the SART (Robinson, Krimsky, et al., 2013) that gave an effect size of 0.56 for the effect of threat of shock on response accuracy to “no-go” distractor stimuli. We wanted 95% power (with alpha 0.05, two tailed) to detect an effect size of 0.56. A power calculation determined that we needed 46 participants. We recruited an extra 4 to allow for ~ 8% participant drop-off. This sample size also has 99% power to detect a reliability of at least 0.5 (the minimum for clinical relevance) at alpha=0.05 (one-tailed). For the final 5-9 month follow up we showed considerable (56%) drop-off. A post hoc matched t-test power analysis showed that with 22 participants (with alpha = 0.05, two tailed) we had only 70.96% power to detect an effect of this magnitude. Notably, however, this still has 83% power to detect a reliability of at least 0.5 (the minimum for clinical relevance) at alpha=0.05 (one-tailed). As such, this three-session analysis is powered for reliability analysis only.

### Stress manipulation

Two electrodes were attached to the back of the participants’ non-dominant wrist. A Digitimer DS5 Constant Current Stimulator (Digitimer Ltd., Welwyn Garden City, UK) delivered the shocks. A short shock-level work up increased the level of the shock until the subject rated it as “unpleasant, but not painful” (Schmitz & Grillon, 2012). As in previous versions of this task (Robinson, Krimsky, et al., 2013) during a threat block, in which the background was red, the participants were told they were at risk of an unpredictable shock (which was independent of their behavioural response). When in a safe block, the background was blue (and participants were told that no shocks would be delivered). Colours were not counterbalanced as prior work has shown this effect to be independent of background colour (C Grillon, Ameli, Merikangas, Woods, & Davis, 1993; Christian Grillon, Baas, Cornwell, & Johnson, 2006).

### Task structure

Participants completed a previously used task (Robinson, Krimsky, et al., 2013) recoded using the Cogent (Wellcome Trust Centre for Neuroimaging and Institute of Cognitive Neuroscience, UCL, London, UK) toolbox for Matlab (2014b, The MathWorks, Inc., Natick, MA, United States). Participants were instructed to respond to “go” target stimuli (“=”) by pressing the space bar as quickly as possible, and withhold a response to “no go” target stimuli (“O”). They were instructed to make their response using their dominant hand. 47 “go” stimuli and 5 “no-go” stimuli were presented in each block. The stimuli were presented for 250ms, followed by an interstimulus interval of 1750ms, before presentation of the next stimulus. There were 8 blocks in total, alternating between threat and safe blocks (order counterbalanced). Each block lasted 104 seconds (See Figure 1). For 3 seconds at the beginning of each block, “YOU ARE NOW SAFE FROM SHOCK!” or “YOU ARE NOW AT RISK OF SHOCK!” appeared on the screen. Participants received a shock in the first threat block, (after trial 45), the second threat block (after trial 8), and the fourth threat block (after trial 17). Total task duration was approximately 14 minutes and 30 seconds [^1^ Script: https://figshare.com/articles/SART_script/3443093].

**Figure 1.**
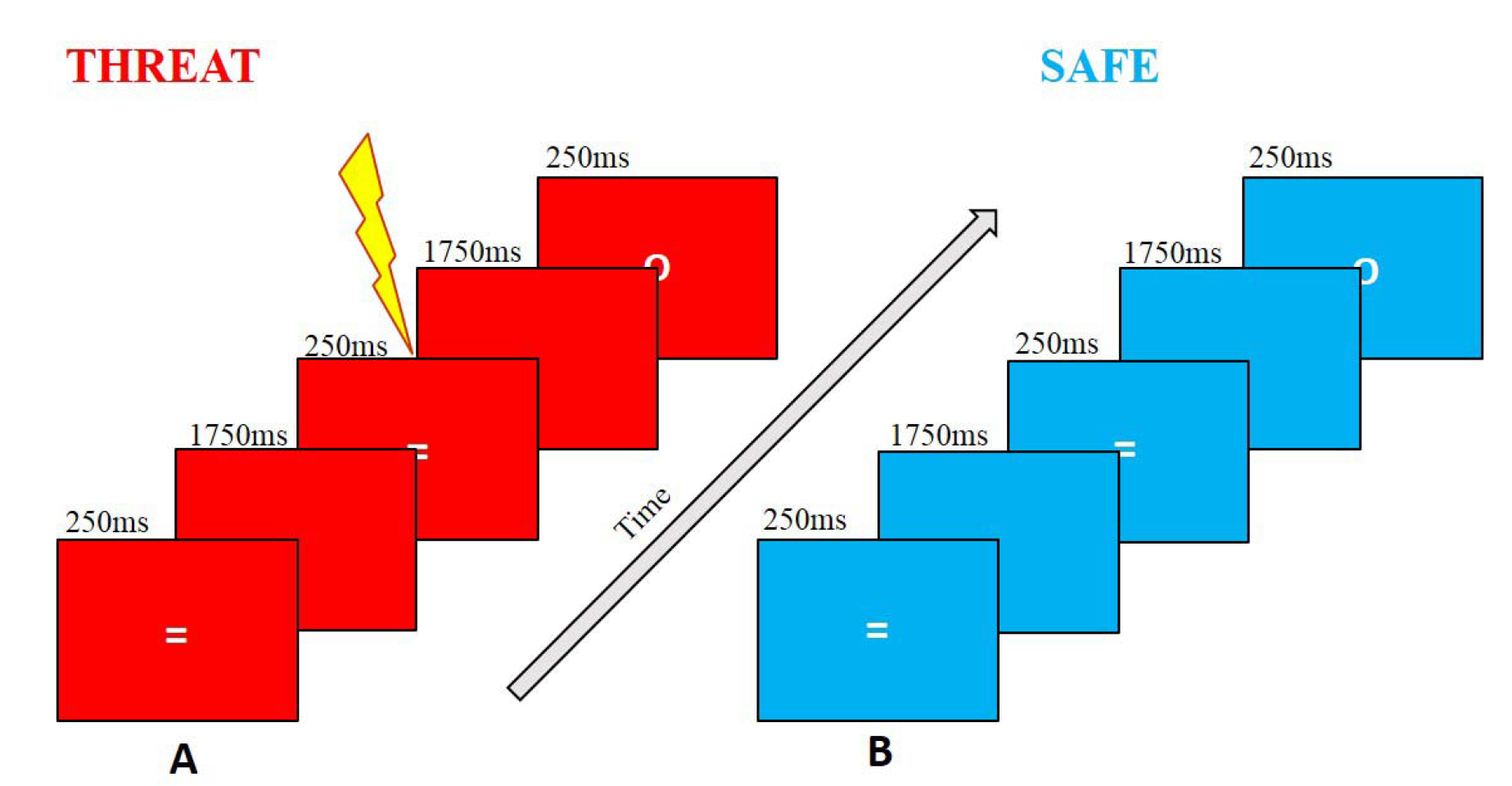
Participants were instructed to press the space bar as quickly as possible for “go” stimuli and withhold responses to infrequent “no-go” stimuli. A: Participants received an unpredictable electric shock (independent of behavioural response) during the threat condition. B: Participants were not at risk of shock during the safe condition.

### Wider sample

Stress-potentiated task performance data from a larger (n=165 heterogeneous sample collected across UCL, UK and NIH, USA are also presented to explore population level statistics.

### Statistical Analyses

Reaction time and accuracy data [^2^ Data: https://dx.doi.org/10.6084/m9.figshare.3398764.v1] were analysed using repeated-measures general linear models in SPSS version 22 (IBM Crop, Armonk, NY). For all analyses, *p* = 0.05, was considered significant. Performance accuracy for each condition (threat/safe) and trial type (“go” / “no-go”) was calculated by dividing the number of correct trials by the total number of trials. As “go” accuracy was 97.5% across two sessions, only “no-go” trials were included in the accuracy analysis. Reaction time analysis was performed on “go” stimuli only as, by definition, “no-go” reaction times are limited and restricted to error trials.

For the first two fully powered sessions, repeated measures ANOVAs were run to investigate reaction time and accuracy differences across conditions. Due to lack of power resulting from attrition (see above) these were not run for the third session.

Task reliability over two and three sessions was tested using two-way mixed model ICCs run in Matlab (2014b) using an “Intraclass Correlation Coefficient” script [^3^ ICC script: (http://uk.mathworks.com/matlabcentral/fileexchange/22099-intraclass-correlation-coefficient--icc-).]. This determined whether the influence of threat of shock on various performance measures remained consistent in individuals between testing sessions. In accordance with (Fleiss, 1986) an ICC coefficient was considered ‘fair to good’ if between 0.4 and 0.75, and ‘excellent’ if above 0.75. In our power calculation we deemed 0.5 the minimum reliability required for clinical relevance. Reliability analyses were completed on the critical delta variables (the difference between threat and safe condition for that variable) to look at the reliability of the *threat-potentiated* effect for reaction time, accuracy and shock rating. Analyses were also run for shock level and trait anxiety scores (for which there are only one measurement per session, so no deltas). Estimates for each condition separately demonstrating the reliability of the individual measures themselves are presented in Table 1 and 2.

## Results

**Table 2:** Reliability of individual measures across two testing sessions (N=50).

| **Individual measures of interest** | **ICC value** | **Repeated measures ANOVA** |
| --- | --- | --- |
| Reaction time to “go” stimuli across safe conditions | 0.82* | $F_{(49,49)} = 5.40$ |
| Reaction to “go” stimuli across threat conditions | 0.91* | $F_{(49,49)} = 11.14$ |
| Accuracy to “no-go” stimuli across safe conditions | 0.82* | $F_{(49,49)} = 5.71$ |
| Accuracy to “no-go” stimuli across threat conditions | 0.87* | $F_{(49,49)} = 7.58$ |
| Self-report anxiety level across safe conditions | 0.63* | $F_{(49,49)} = 2.78$ |
| Self-report anxiety level across threat conditions | 0.62* | $F_{(49,49)} = 2.59$ |

\* $p < 0.05$

**Table 3:**
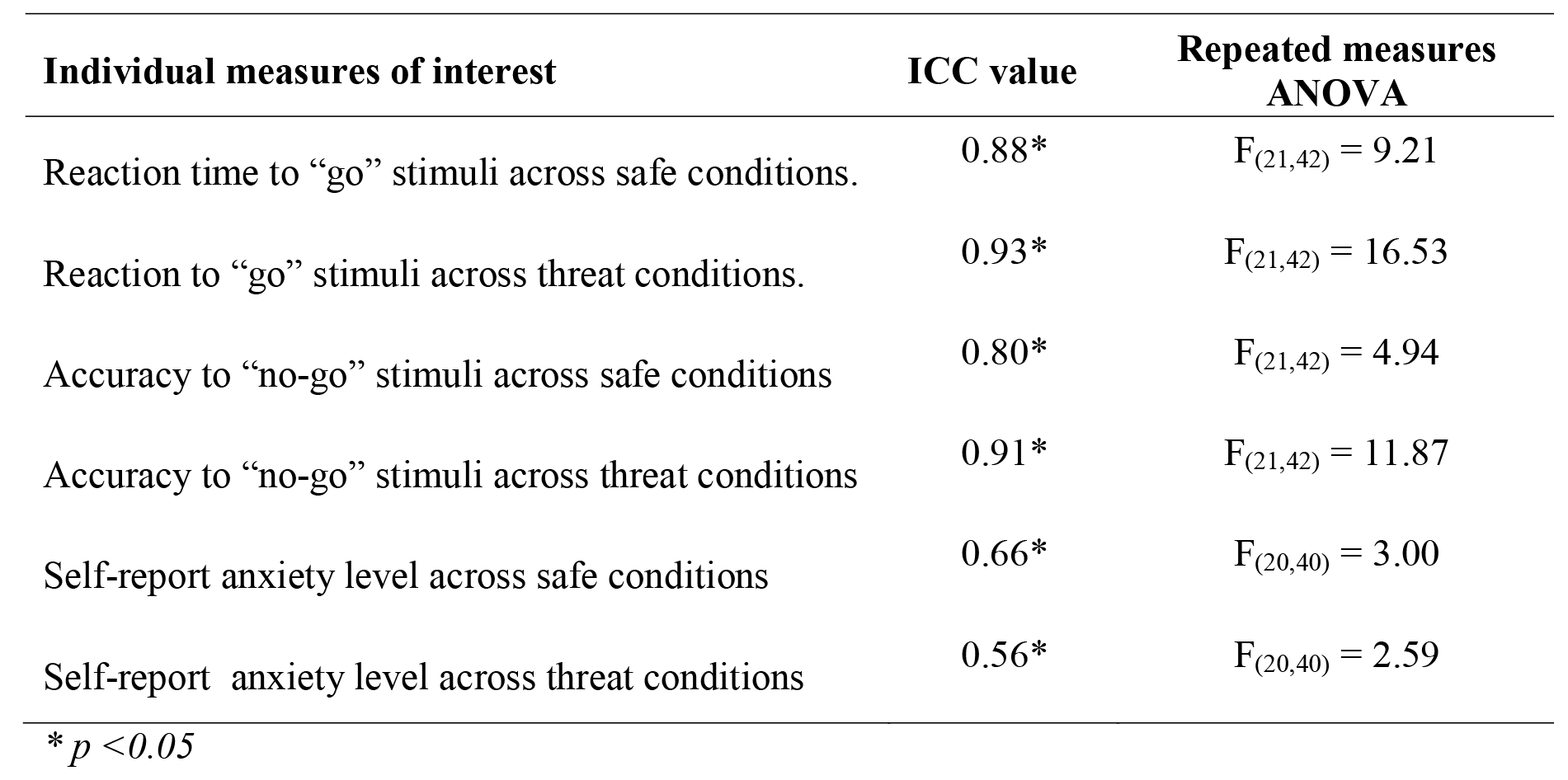
Reliability of individual measures across three testing sessions (N=22).

### Reaction time to “go” stimuli

A repeated measures ANOVA including condition and session revealed a significant effect of condition (F_(1,49)_ = 6.75, *p* = 0.012, 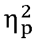 = 0.121) (Figure 1c; 1d). Participants were slower to respond during the threat condition relative to the safe condition (threat mean = 365.40, SD = 64.71; safe mean = 355.44, SD = 56.73). There was no effect of session, nor a session × condition interaction (*ps* > 0.250). Reliability for the effect of threat of shock across 2 sessions was significant and “fair to good” with an ICC of 0.58 (F_(49,49)_ = 2.37, *p* = 0.0015, 95% CI 0.26, 0.76) and remained “fair to good” across 3 sessions, with an ICC of 0.50 (F_(21,42)_ = 1.99, *p* = 0.029, 95% CI -0.017, 0.78).

### Accuracy to “no go” stimuli

A repeated measures ANOVA including condition and session revealed a main effect of condition (F_(1,49)_ = 9.11, *p* = 0.004, 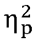= 0.157) (Figure 1a; 1b). Participants were significantly more accurate under threat of shock (threat mean = 0.685, SD = 0.20; safe mean = 0.641, SD = 0.18). There was no main effect of session or a significant threat × session interaction (*p* = 0.93, *p* = 0.063, respectively). The reliability of threat induced accuracy changes across 2 sessions had a non-significant ICC of 0.23 (F_(49,49)_ = 1.31, *p* = 0.17, 95% CI -0.32, 0.55), but across 3 sessions, the effect of threat on accuracy was significant and “fair to good”, with an ICC of 0.51 (F_(21,42)_ = 2.02, *p* = 0.026, 95% CI -0.0026, 0.78).

### Anxiety Rating

Participants were more anxious during the threat condition relative to safe condition (F_(1,49)_ = 225.96, *p* < 0.001, 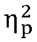 = 0.82)(Figure 1e; 1f). There was no main effect of session or a significant session x condition interaction. The significant reliability of this effect was “fair to good” across 2 sessions with an ICC of 0.66 (F_(49,49)_ = 3.05, *p* < 0.001, 95% CI 0.41, 0.81) and across 3 sessions with an ICC of 0.74 (F_(20,40)_ = 3.90, *p* < 0.001, 95% CI 0.48, 0.89).

### Shock Level

Shock level was significantly higher in the second session (F_(1,49)_ = 5.96, *p* = 0.018, 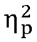 = 0.108; session 1 mean = 6.35, SD = 3.15; session 2 mean = 7.12, SD = 2.81)(Figure 1g). Reliability for shock level over 2 sessions was not significant, with an ICC of 0.27 (F_(49,49)_ = 1.37, *p* = 0.13, 95% CI -0.28, 0.59) and across 3 sessions with an ICC of 0.29 (F_(21, 42)_ = 1.40, *p* = 0.17, 95% CI, -0.46, 0.68).

### Trait Anxiety

Trait anxiety scores were also analysed as a comparison. There was no significant change over session (*p* = 0.622) (Figure 1h) and the reliability of the trait anxiety score was “excellent”, with an ICC of 0.95 across 2 sessions (F_(49,49)_ = 20.97, *p* < 0.001, 95% CI 0.92, 0.97) and across 3 sessions 0.94 (F_(21,42)_ = 16.67, *p* < 0.001, 95% CI, 0.87, 0.97).

### Population variance

Accuracy to “no go” stimuli in 165 subjects revealed a population mean of 0.072, median of 0.05, a standard deviation of 0.15 (Figure 1).

**Figure 2:**
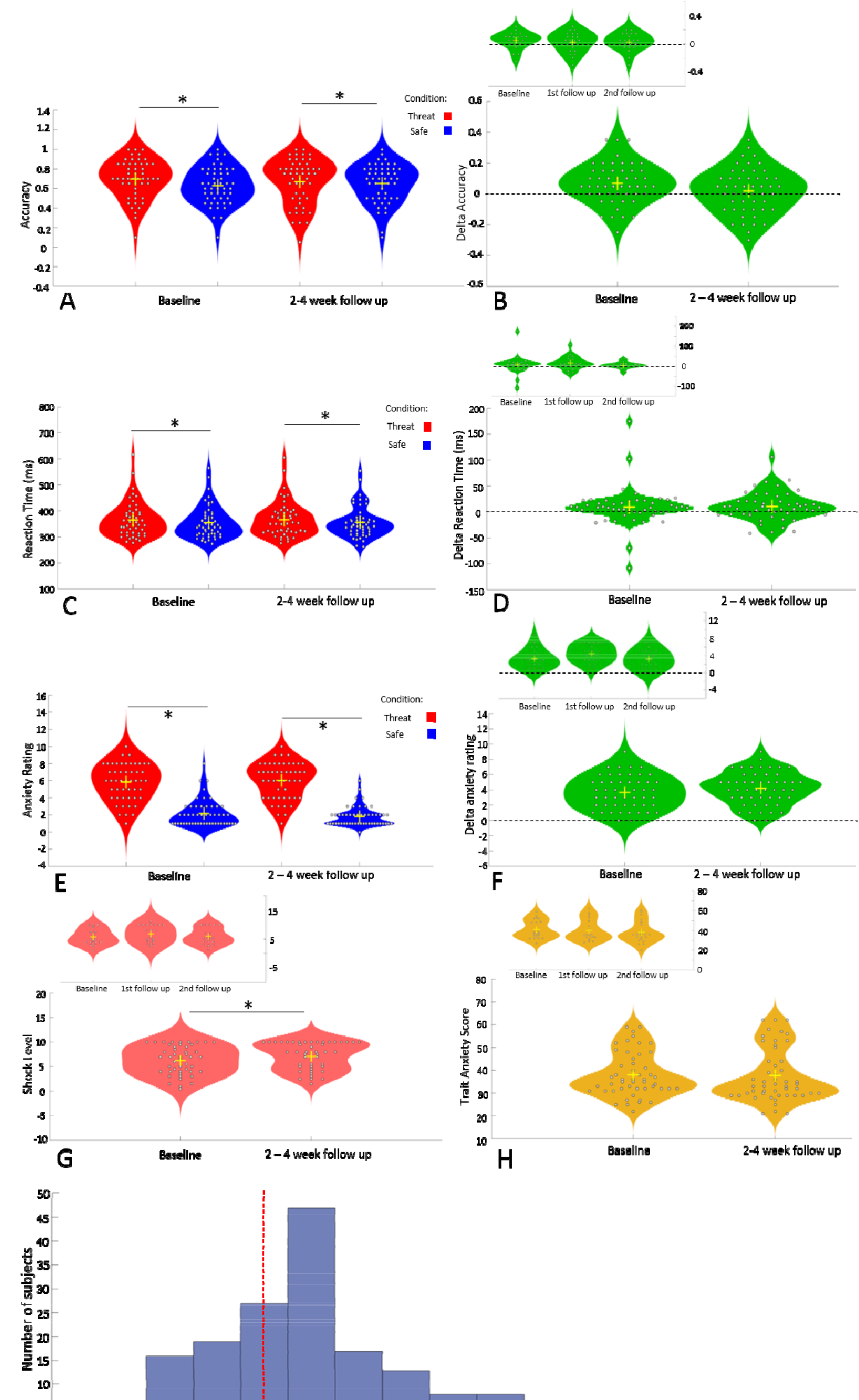
Violin plots (shaded area represents a histogram) **A**. Reaction time to “go” stimuli across threat and safe conditions. There was a main effect of condition (*p* = 0.012). **B**. Delta reaction time (inset across 3 sessions). **C**. Accuracy to “no go” stimuli across threat and safe conditions. There was a main effect of condition (*p* = 0.04) **D**. Delta accuracy (inset across three sessions). **E**. Anxiety rating across threat and safe conditions. There was a main effect of condition (*p* = 0.05). **F**. Delta anxiety ratings (inset across three conditions). **G**. Shock level across baseline and follow up (main effect of session *p* = 0.018; inset shock level across three sessions). **H**. Trait anxiety score across testing sessions (inset trait anxiety across three sessions). **I**. Distribution of delta distractor accuracy scores on the SART in a large population (N = 165). Dotted line at zero demonstrates population as a whole shifted towards threat-potentiated accuracy.

## Discussion

We show that threat of shock can reliably shift within-subject cognitive and self-report measures of stress across three sessions and three quarters of a year, improving accuracy to distractor stimuli and slowing down responses to target stimuli.

Importantly, the impact of stress on this task also shows good within-subject reliability over 2–4 weeks and again over a 5-9 months. For reference, this means that the emotional manipulation (i.e. threat of shock) on this task, is considerably more reliable than the emotional manipulation (i.e. emotional stimuli) on the emotional Stroop and dot probe tasks (Adams et al., 2015; Kindt et al., 1996)(see Table 1).

Stress tests are of great value in cardiac medicine as they are able to identify patients who may be more vulnerable and need closer monitoring around surgery, which in turn leads to improved outcomes (Wijeysundera, Beattie, Austin, Hux, & Laupacis, 2010). According to the diathesis model of mood disorders, when stressful life events are coupled with an underlying vulnerability to mood disorders and a threshold is reached, a disorder is triggered. Understanding individual responses to stress is therefore key to understanding the differences in vulnerability of mood and anxiety disorders. Depression and anxiety constitute some of the most common psychiatric disorders, and it is suggested that over the next 20 years these rates will continue to rise (WHO, 2013). There are poor treatment outcomes in depression for pharmacological and therapeutic approaches (Trivedi et al., 2006; Cuijpers et al., 2014). Identifying those who would benefit from particular treatments (Kazdin, 2007), or even vulnerability prior to disorder onset with a non-subjective cognitive task could consequently lower costs and reduce time in treatment. Additionally, cognitive paradigms which show good reliability are important for research, impacting replicability and the accurate interpretation of existing findings (Open Science Collaboration, 2015).

It should be noted that self-report trait anxiety also has a high ICC in this study. However, interpretation of this is limited due to anchoring effects (Tversky & Kahneman, 1974) and demand characteristics (Weber & Cook, 1972). Our task is not obviously subject to these effects and also benefits from being a concurrent (i.e. not retrospective) measure. The poorer test retest reliability on our accuracy delta variable for the two to four week follow up may be due to a reduction in power resulting from the smaller number of no-go responses, and suggests that go reaction time differences may prove the more reliable target.

In summary, we argue that the impact of threat of shock on cognition might hold promise as a putative stress test for emotional responding.

## Acknowledgements

This research was funded by a Medical Research Foundation Equipment Competition Grant (C0497, Principal Investigator OJR), and a Medical Research Council Career Development Award to OJR (MR/K024280/1). OJR has received consulting fees from IESO digital health unrelated to the current project.

We thank Jonathan P. Roiser for his advice and helpful comments on the manuscript.

This manuscript has open data and open materials

